# Number-Space Association in Macaques

**DOI:** 10.64898/2026.03.23.713206

**Authors:** G. Annicchiarico, M. Belluardo, G. Vallortigara, P.F. Ferrari

## Abstract

Humans order numbers in space from left to right, with smaller quantities represented preferentially in the left hemispace and larger ones in the right hemispace. The direction of this mental number line (MNL), or more generally of number–space associations (NSA), is influenced by cultural habits such as reading and writing direction. However, a growing body of evidence from pre-verbal infants and non-human animals suggests that number–space mappings may also have biological foundations. In non-human primates, evidence for a directional MNL remains mixed, partly due to small sample sizes and methodological heterogeneity. Here, we tested samples of rhesus (*Macaca mulatta)* and crab-eating macaques (*Macaca fascicularis*) across two experiments using spontaneous food-related tasks. In Experiment 1, monkeys chose between identical food quantities (1×1 to 24×24) presented on the left and right. No systematic spatial choice bias emerged as a function of numerical magnitude, and hand use did not differ across exact numerical pairs, although exploratory analyses revealed magnitude-related modulations of manual responses. In Experiment 2, monkeys were habituated to small (4×4) or large (16×16) quantities and subsequently tested with the alternative quantity. Result showed significantly more leftward choices following numerical decreases (16→4) and more rightward choices following numerical increases (4→16), indicating that relative numerical context, rather than absolute magnitude, elicited directional spatial biases. These findings suggest that in macaques, number–space associations emerge most robustly in comparative contexts involving expectancy violations of magnitude.

## Introduction

Numerical magnitudes in humans are known to be organized along a spatial continuum referred to as the Mental Number Line (MNL), with smaller numbers on the left and larger ones on the right (Galton, 1880; Dehaene, 1992). Dehaene, Bossini, and Giraux (1993) showed that people respond more quickly to small numbers with left-hand responses and to large numbers with right-hand responses, a phenomenon dubbed the Spatial-Numerical Association of Response Codes (SNARC) effect. Early studies suggested that the orientation of the Mental Number Line (MNL) is caused by cultural practices—such as reading and writing direction. For example, individuals who read from right to left, like Arabic speakers, exhibit a reversed spatial-numerical mapping (Zebian, 2005), whereas those exposed to bidirectional reading show no consistent SNARC effect (Shaki et al., 2009). These findings support the idea that MNL orientation is culturally constructed (see also Pitt et al., 2021). However, a growing body of evidence from preverbal infants and non-human species points instead to a biological and evolutionary basis for spatial-numerical associations (Adachi, 2014; Bulf et al., 2016; De Hevia et al., 2014; Di Giorgio et al., 2019; Drucker & Brannon, 2014; Lourenco & Longo, 2010; Rugani et al., 2014, 2016; Vallortigara, 2018). For instance, de Hevia et al. (2017) habituated 0–3-day-old neonates to a long line with many sounds or a short line with few sounds. When subsequently shown two lines on either side, infants looked longer to the left for shorter/fewer stimuli and to the right for longer/more numerous stimuli. In a similar vein, Di Giorgio et al. (2019) tested newborns (∼51 h old) using habituation to specific numerosities, and found that they looked left for smaller and right for larger numerosities. Additional support for a biologically grounded MNL comes from studies on birds (Rugani et al., 2007, 2010, 2015), non-human primates (Adachi, 2014; Drucker & Brannon, 2014; Gazes et al., 2017; Johnson-Ulrich & Vonk, 2018; Rugani et al., 2024), and even insects (Giurfa et al., 2022), suggesting that spatial-numerical associations may reflect an evolutionarily conserved cognitive mechanism.

A key demonstration of a bidirectional MNL comes from Rugani et al. (2015), who showed that three-day-old chicks trained to associate a panel with five dots to a food reward consistently chose the left panel for smaller numbers (e.g., 2) and the right for larger ones (e.g., 8). When trained with a larger reference number (20 dots), the same chicks selected the left panel for 8 and the right for 32, indicating that their choices were based on relative magnitude—a core feature of the MNL. Similar findings have been replicated in bees (Giurfa, 2022) and in human newborns (Di Giorgio, 2019). In non-human primates, spatial-numerical associations have also been observed: gorillas, orangutans (Gazes et al., 2017), and American black bears (Johnson-Ulrich & Vonk, 2018) exhibit SNARC-like effects, though individual variation in directionality occurs. Chimpanzees trained to select Arabic numbers in ascending order responded faster when “1” appeared on the left and “9” on the right (Adachi, 2014), although this may reflect learned sequential patterns rather than spontaneous magnitude processing.

Macaques represent an ideal model for investigating number–space associations due to their close social, cognitive, emotional, physiological, and genetic similarities to humans (Phillips et al., 2014; Rosati et al., 2016; Sallet, 2022; Wang et al., 2020; Xue et al., 2016). Behavioral evidence demonstrates that macaques can discriminate numerosities. (Cantlon & Brannon, 2006; Merten & Nieder, 2009; Nieder, 2012). Neurophysiological studies have identified “number neurons” in frontal, parietal, and temporal cortices that selectively respond to specific numerosities (Nieder, 2016; Nieder et al., 2002; Nieder & Miller, 2003; Okuyama et al., 2015; Viswanathan & Nieder, 2015), and the intraparietal sulcus (IPS) shows overlapping neural substrates for numerical and spatial processing, paralleling findings in humans (Hubbard et al., 2005), crows (Ditz & Nieder, 2015), chicks (Kobylkov et al. 2022) and zebrafish (Luu et al., 2024).

Behavioral studies in macaques suggest a spatial mapping of numbers (Beran et al., 2019; Drucker & Brannon, 2014; Rugani et al., 2024) but the existence of a consistent left-to-right MNL remains uncertain. Drucker and Brannon (2014) found that monkeys trained to select the fourth dot from the bottom in a vertical array chose the fourth from the left in a horizontal layout, possibly indicating a left-to-right bias—though this could also reflect a right-to-left strategy (i.e., second from the right), leaving the directionality ambiguous. A recent replication using the same method, however, failed to reproduce this effect, as monkeys did not reliably choose the fourth position from the left over the fourth from the right (Silva et al., 2025). In another study, Rugani et al. (2024) reported a left-to-right mapping in two rhesus macaques asked to recall target positions within arrays of 2 to 10 items. Accuracy was higher on the left for smaller numerosities and shifted rightward with increasing quantities, yet this pattern was inconsistent across conditions and limited by a small sample size. In contrast, Beran et al. (2019) adapted Rugani et al.’s (2015) two-choice paradigm to a touchscreen task, where monkeys habituated to a central reference numerosity were then presented with smaller and larger quantities on the left and right. The authors did not find any consistent directional bias at the group and individual level. However, monkeys were subsequently required to choose between two distinct arrays of dots presented simultaneously on either side, 12 out of 19 subjects showed individual spatial biases—some favoring smaller numerosities on the left, others on the right.

Given these inconclusive and often contradictory findings, here we designed two food-based tasks to test whether macaques spontaneously map numerical magnitudes onto space. Food was selected as an ecologically relevant stimulus, as numerical abilities in non-human animals likely evolved in foraging contexts, and food-based designs are more likely to elicit automatic behaviors. Consistent with Eccher et al. (2023), the SNARC effect tends to emerge in implicit tasks without explicit numerical judgments. Similarly, capuchins discriminate food quantities more efficiently than tokens, due to the higher motivational value of food (Addessi et al., 2008). Building on this rationale, we conducted two spontaneous food-based tasks in rhesus and crab-eating macaques to examine whether spatial–numerical mappings emerge under ecologically relevant and implicitly structured conditions.

In the first experiment, we investigated spontaneous number–space associations by presenting macaques with equal numerical quantities simultaneously on the left and right sides, ranging from 1 to 24, without any reference number or prior habituation. This design aimed to reveal potential inherent spatial biases, assessing whether monkeys naturally associate smaller or larger numerosities with the left or right hemispace, respectively.

The second experiment employed a habituation-test paradigm involving two specific numerosities. During habituation, subjects were repeatedly presented with a reference quantity to establish a baseline. In the test phase, they were shown either a larger or smaller numerosity, allowing us to examine relative spatial-numerical biases. This approach—first used by Rugani et al. (2015) with chicks and later adapted for bees (Giurfa et al., 2022) and human newborns (Di Giorgio et al., 2019)—consistently revealed a left-to-right spatial mapping in these species. Beran et al. (2019) implemented a computerized version of this paradigm in macaques but did not observe the same pattern.

In our replication, we introduced key modifications to enhance ecological validity. Most notably, we used real food items as stimuli to increase task salience and motivation, based on evidence that food-related numerosity elicits more robust responses in non-human primates (Addessi et al., 2008; Beran et al., 2005). Moreover, unlike previous studies using centrally presented stimuli (Di Giorgio et al., 2019; Giurfa et al., 2022; Rugani et al., 2015), our design allowed for lateralized choices during habituation trials. This methodological shift enabled a more direct comparison between habituation and test responses, allowing us to trace potential changes in spatial preferences as a function of numerical deviation.

## Materials and Methods

### Experiment 1

#### Subjects

We tested 17 macaques, 7 rhesus macaques (*Macaca mulatta*) and 10 crab-eating macaques (*Macaca fascicularis*), housed at the Institut des Sciences Cognitives Marc Jeannerod (Lyon, France). The age of the monkeys ranged from 3 to 14 years (mean age = 5.38; sd = 3.32) (see Table 1). The subjects were housed in colony cages with one or more partners. During the test they were isolated in a testing cage (220 cm x 70 cm X 115 cm) and they were free to move without any hindrance. Fresh fruit, pellet and nuts were provided daily. Water was available ad libitum.

**Table 1.**
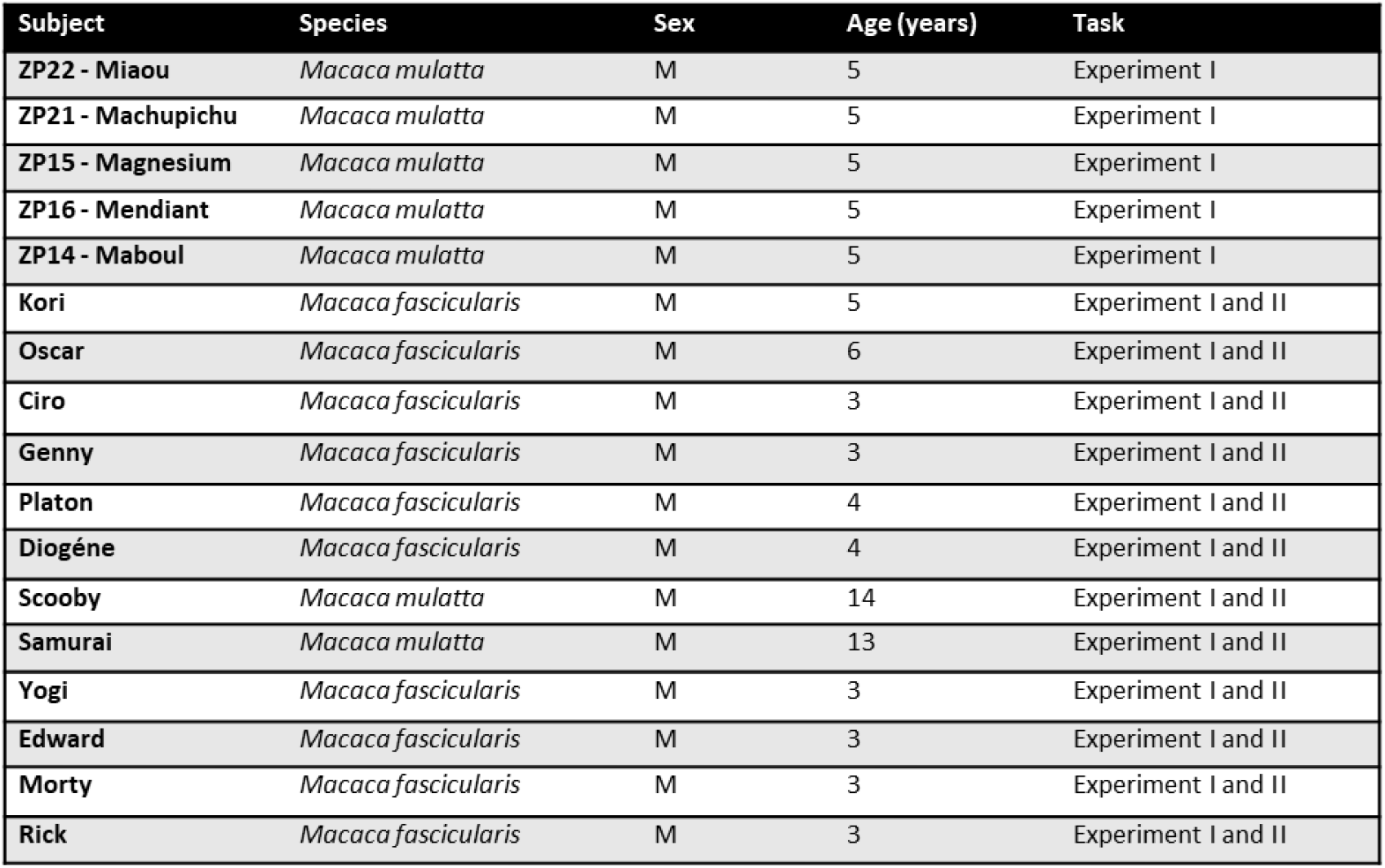
Table of information (Names, Species, Sex, Age) of experimental subjects in Experiment I and/or II.

| Subject | Species | Sex | Age (years) | Task |
| --- | --- | --- | --- | --- |
| ZP22 - Miaou | *Macaca mulatta* | M | 5 | Experiment I |
| ZP21 - Machupichu | *Macaca mulatta* | M | 5 | Experiment I |
| ZP15 - Magnesium | *Macaca mulatta* | M | 5 | Experiment I |
| ZP16 - Mendiant | *Macaca mulatta* | M | 5 | Experiment I |
| ZP14 - Maboul | *Macaca mulatta* | M | 5 | Experiment I |
| Kori | *Macaca fascicularis* | M | 5 | Experiment I and II |
| Oscar | *Macaca fascicularis* | M | 6 | Experiment I and II |
| Ciro | *Macaca fascicularis* | M | 3 | Experiment I and II |
| Genny | *Macaca fascicularis* | M | 3 | Experiment I and II |
| Platon | *Macaca fascicularis* | M | 4 | Experiment I and II |
| Diogène | *Macaca fascicularis* | M | 4 | Experiment I and II |
| Scooby | *Macaca mulatta* | M | 14 | Experiment I and II |
| Samurai | *Macaca mulatta* | M | 13 | Experiment I and II |
| Yogi | *Macaca fascicularis* | M | 3 | Experiment I and II |
| Edward | *Macaca fascicularis* | M | 3 | Experiment I and II |
| Morty | *Macaca fascicularis* | M | 3 | Experiment I and II |
| Rick | *Macaca fascicularis* | M | 3 | Experiment I and II |

All housing and procedures used in this research complied with the current guidelines in the matter of care and use of laboratory animals (European Community Council Directive No. 86–609), and were authorized by our local ethics board (03.10.18) and the French Ministry of Research (10.10.18; authorization no. 15091_2018071014483295_v2).

#### Apparatus

The apparatus consisted of a testing cage with a frontal PVC plexiglass panel (122 cm x 46 cm x 3 cm) (Fig.1). The PVC panel had two holes at the bottom, one on the left and one on the right, each positioned 10.5 cm from the border. A shelf (25 cm x 83 cm x 1 cm) was attached to the cage to allow the experimenter to place two food containers in front of the monkey. The food containers comprised two test tube holders (8 cm x 12.5 cm x 2 cm), positioned on the left and right parts of a tray (a wooden support measuring 12 cm x 38 cm x 2 cm), equidistant from the board. Each test tube holder contained 24 holes (diameter 1.8 cm), spaced 2 mm apart. A metal stick was attached to the shelf to designate the point from which the monkeys could reach the food. This spatial index was used by the experimenter to place the tray within the monkey’s “reachable space”.

**Figure 1.**
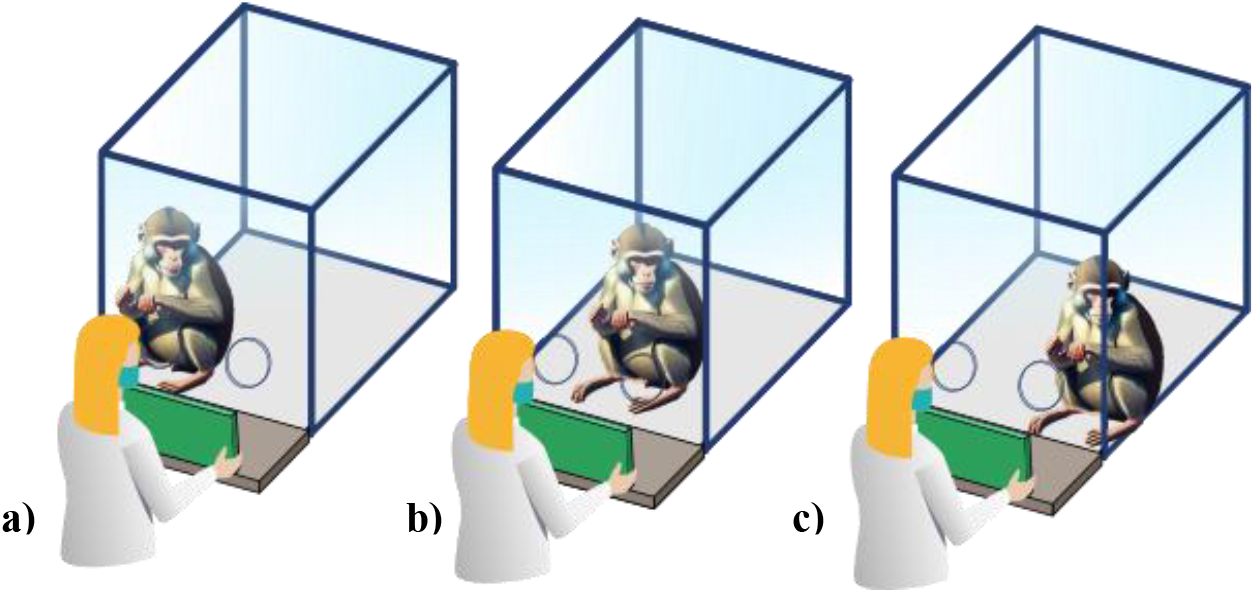
Illustration of stimulus presentation with the monkey positioned to the right (a), center (b), or left (c) relative to the stimuli.

#### Experimental Procedure

The experimental procedure was adapted from Rugani et al (2015), with several modifications. We designed an equal comparison quantity task where numerical targets were represented by pieces of food instead of dots on a panel, ensuring the stimuli were ecologically relevant to the monkeys. We chose to exclude a training phase with a reference number in order to observe the subjects’ spontaneous behavioral responses to varying quantities within a numerical range.

Each monkey was pseudorandomly presented with six possible pair-wise combinations of the same quantity of food pieces in the left and right food boards: 1 × 1, 2 ×. 2, 4 × 4, 8 × 8, 16 × 16 and 24 × 24. The food pieces were raisins, placed by the experimenter inside the holes of both tube racks (Fig. 2a).

**Figure 2.**
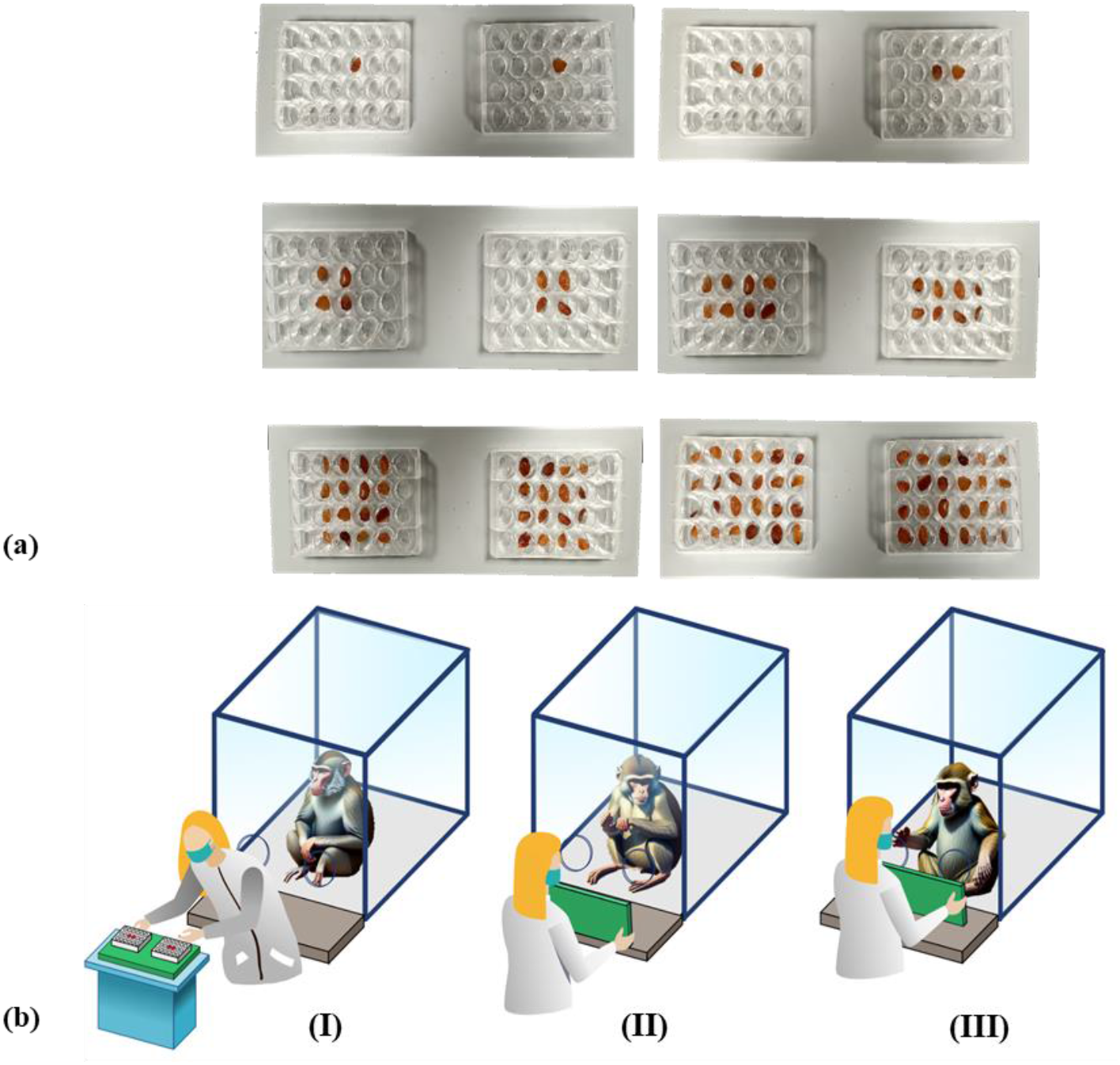
(a) The six possible pair-wise combinations of the same quantity of pieces of food placed in the left and in the right food container. (b) Illustration of the experimental procedure: (I) Experimenter prepares food containers with back turned to monkey; (II) Experimenter shows tray out of reach, waiting for monkey to look; (III) Experimenter brings containers within reach, monkey chooses left/right and takes one raisin.

For each trial, the experimenter arranged the food containers with her back turned to the monkey, preventing the subject from viewing the contents before the trial began. The complete experimental procedure is illustrated step-by-step in Figure 2b.

If the monkey did not direct its gaze towards both food containers within ten seconds, the trial was terminated, and the tube racks were removed. Each subject underwent an experimental session consisting of thirty trials, with each of the six food pairwise combinations presented five times in a pseudo-random order.

As soon as the subject chose, reached, and grasped one raisin from one of the two (left or right) food containers, the experimenter removed the containers. Limiting the reward to only one piece of food per trial was intended to prevent satiation and ensure that the subject maintained sufficient motivation to continue and complete the experiment. If the monkey did not take any food within ten seconds, the experimenter removed the tube racks.

The procedure was repeated for 30 subsequent successful trials, defined as trials where the monkey looked at both food containers and reached for a piece of food from one of them. The subjects did not undergo any training sessions but only a familiarization phase to become comfortable with the experimental apparatus. Each subject was tested individually and isolated from the group during the experiment.

All experimental sessions have been video recorded through one camera (Digital Videocamera Panasonic HC V-180 Full-HD optical zoom 90×), allowing offline data elaboration.

#### Data Analyses

For each animal, the experimental session was coded offline using Avidemux 2.7.8 Release by the experimenter. A second researcher coded a random 20% of videos to assess reliability. The reliability scores between coders were very high (all k > 0.8). For each experimental trial, the following variables were coded: the number of stimuli presented to the monkey (1, 2, 4, 8, 16 or 24) the choice made by the subject (right or left food container), the hand used to grasp the food, the quality of the trials (1 if the experimenter was uncertain whether the monkey had truly attended to both food containers or if the monkey showed stereotyped behavior such as repeated swapping ; 2 if the monkey was not completely centered and/or had a hand resting on/near one of the plexiglass holes; 3 if the monkey was centrally positioned with no visible hand bias or other confounds), and the position of the monkey relative to the two holes (2 if the monkey was positioned too far to the right; 0 if too far to the left; 1 if centrally aligned). One subject was excluded from the analysis due to its biased choice (right choices) throughout the entire task.

The six possible pair-wise combinations of the same quantity of pieces of food (1 × 1, 2 × 2, 4 × 4, 8 × 8, 16 × 16, 24 × 24) were classified as follows: 1 × 1, 2 × 2, 4 × 4, and 8 × 8 were considered “small quantities,” whereas 16 × 16 and 24 × 24 were defined as “large quantities.”

## Statistical analyses and Results

### Choice Preference Analysis

#### Statistical Analysis of Choice as a Function of Stimulus Pair Identity

To evaluate whether choice behavior was influenced by potential nuisance variables, we first conducted a series of control analyses. Trial Quality and Subject Position (left/center/right) were each tested independently using generalized linear mixed-effects models (GLMMs) with a binomial distribution and a logit link function. All models included a random intercept for Subject ID to account for repeated measures.

Type III Wald χ^2^ tests (car package) were used to evaluate the significance of fixed effects. Results indicated that Subject Position (χ^2^(2)=25.29, p*** < .001) significantly predicted choice behavior, whereas Trial Quality (χ^2^(2)=4.35, p = .114) did not (Table S1). Therefore, Position was retained as covariate in all subsequent analyses.

To assess whether monkeys’ spatial choices varied as a function of the numerical pair presented, we fitted a GLMM with a binomial distribution and logit link function. The binary dependent variable was the subject’s choice (0 = left, 1 = right). The main predictor of interest was the numerical pair identity (Pair), corresponding to the six numerical quantities used in the experiment (1, 2, 4, 8, 16, and 24 items).

Based on the preliminary analyses, Subject Position was included as fixed-effect covariates. A random intercept for Subject ID was included to account for repeated trials within individuals. Model estimation was performed using the lme4 package (Bates et al., 2015) in R 4.3.1 (R Core Team, 2023). Fixed effects were evaluated using Type III Wald χ^2^ tests from the car package (Fox & Weisberg, 2018).

### Results

The GLMM revealed a significant effect of Position (*χ*^2^(2) = 22.72, *p\*\*\** < .001), indicating that the spatial location of the monkey significantly influenced choice behavior. However, the effect of Pair, representing the numerical quantity presented, was not statistically significant (*χ*^2^(5) = 6.44, *p* = .266), suggesting that monkeys did not systematically choose one side over the other based on the specific numerical pairing (Table S2 ; Fig.3).

**Figure 3.**
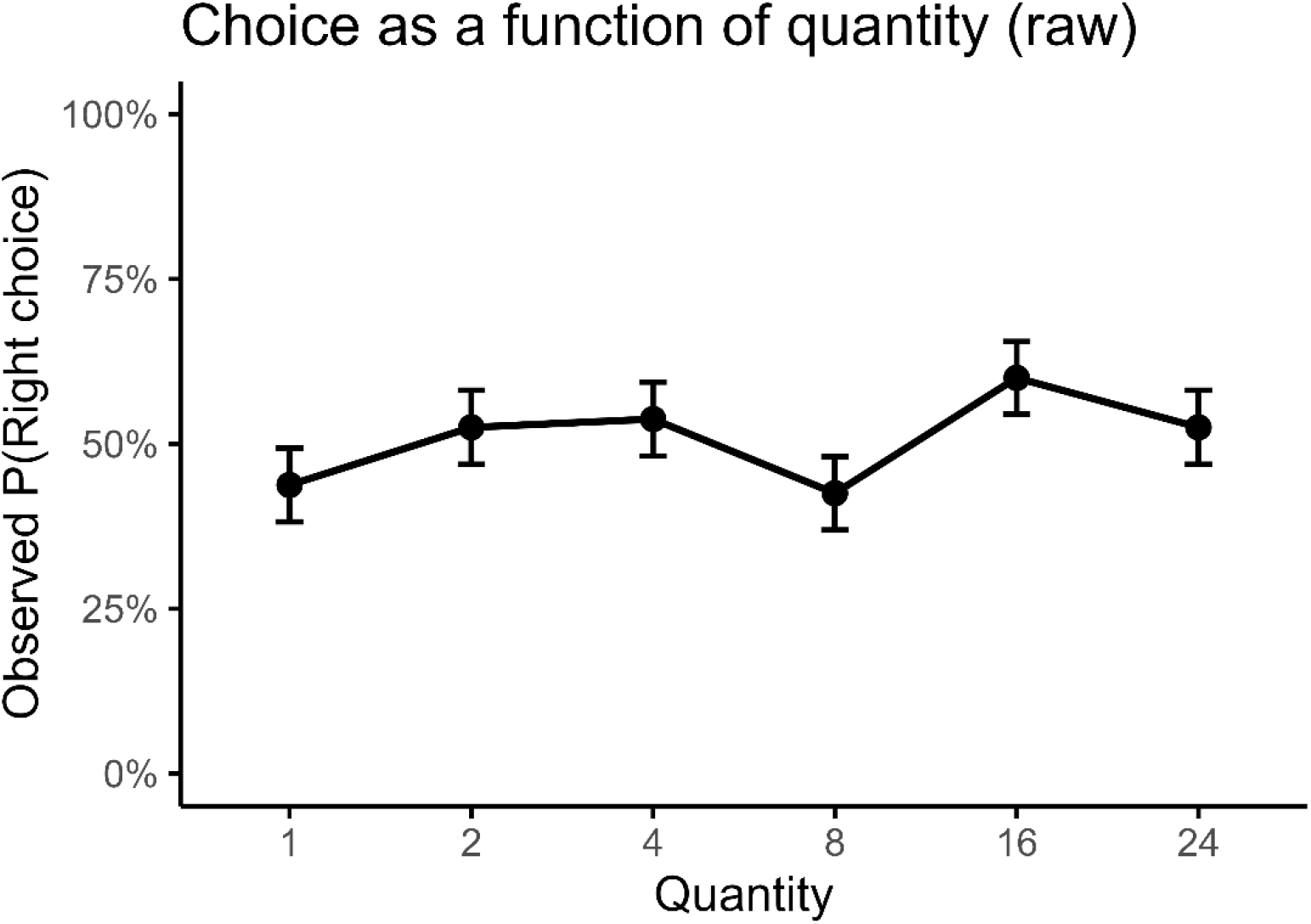
Choice performance across numerical amounts (raw proportions). Mean proportion of right-side choices across the six numerical amounts (1, 2, 4, 8, 16, 24 items). Dots represent group means and vertical bars indicate ±1 standard error (SE). No systematic lateral bias as a function of quantity was observed.

### Statistical Analysis of Choice as a Function of Numerical Magnitude

Given the wide numerical range tested in Experiment 1, we conducted additional exploratory analyses to assess whether choice behavior was modulated by numerical magnitude treated as a continuous or coarse-grained variable. First, we fitted a GLMM including a log2-transformed and centered numerical predictor, while controlling for Subject Position. The logarithmic linear trend associated with numerical magnitude was not statistically significant (χ^2^(1) = 2.08, p = .149; Table S3). A likelihood ratio test comparing the full model with a reduced model excluding numerical magnitude also failed to show a significant improvement in model fit (Δχ^2^(1) = 2.07, p = .151; Table S4).

In a complementary analysis, numerical pairs were regrouped into two categories reflecting overall magnitude (small: 1–8 items; large: 16–24 items). A GLMM including Amount_group (Small/Large) as a fixed effect and Subject Position as a covariate revealed no significant effect of numerical grouping on choice behavior (χ^2^(1) = 2.26, p = .132; Table S5; Fig. S1), while Position remained highly significant (χ^2^(2) = 24.05, p < .001).

Taken together, neither continuous nor categorical representations of numerical magnitude supported a systematic effect on spatial choice in Experiment 1.

### Hand Preference Analysis

#### Statistical Analysis of Hand as a Function of Stimulus Pair Identity

Before assessing the effect of numerical quantity on hand use, we first evaluated potential nuisance predictors. Trial Quality and Subject Position (left/center/right) were each tested separately using generalized linear mixed-effects models (GLMMs) with a binomial distribution and logit link function. All models included a random intercept for Subject ID to account for repeated measures. Significance of fixed effects was assessed using Type III Wald χ^2^ tests.

Control analyses revealed that none of the factors significantly influenced hand choice (Table S6). Accordingly, they were not retained as covariates in all subsequent models.

To determine whether hand preference varied as a function of the numerical quantity displayed, we fitted a GLMM with a binomial distribution and logit link function. The dependent variable was the hand used (0 = left, 1 = right), and the primary predictor was numerical pair identity (Pair), corresponding to the six quantities used in the experiment (1, 2, 4, 8, 16, and 24 items). A random intercept for Subject ID accounted for repeated trials within individuals. Model estimation was performed using the *lme4* package, and Type III Wald χ^2^ tests (car package) were used to evaluate fixed effects.

#### Results

The analysis revealed no significant effect of Pair on hand preference (χ^2^(5) = 9.903, *p* = .078), indicating that monkeys did not systematically use one hand more than the other as a function of numerical magnitude. Thus, hand preference did not reliably vary across the six numerical quantities tested (Table S7; Fig. 4).

**Figure 4.**
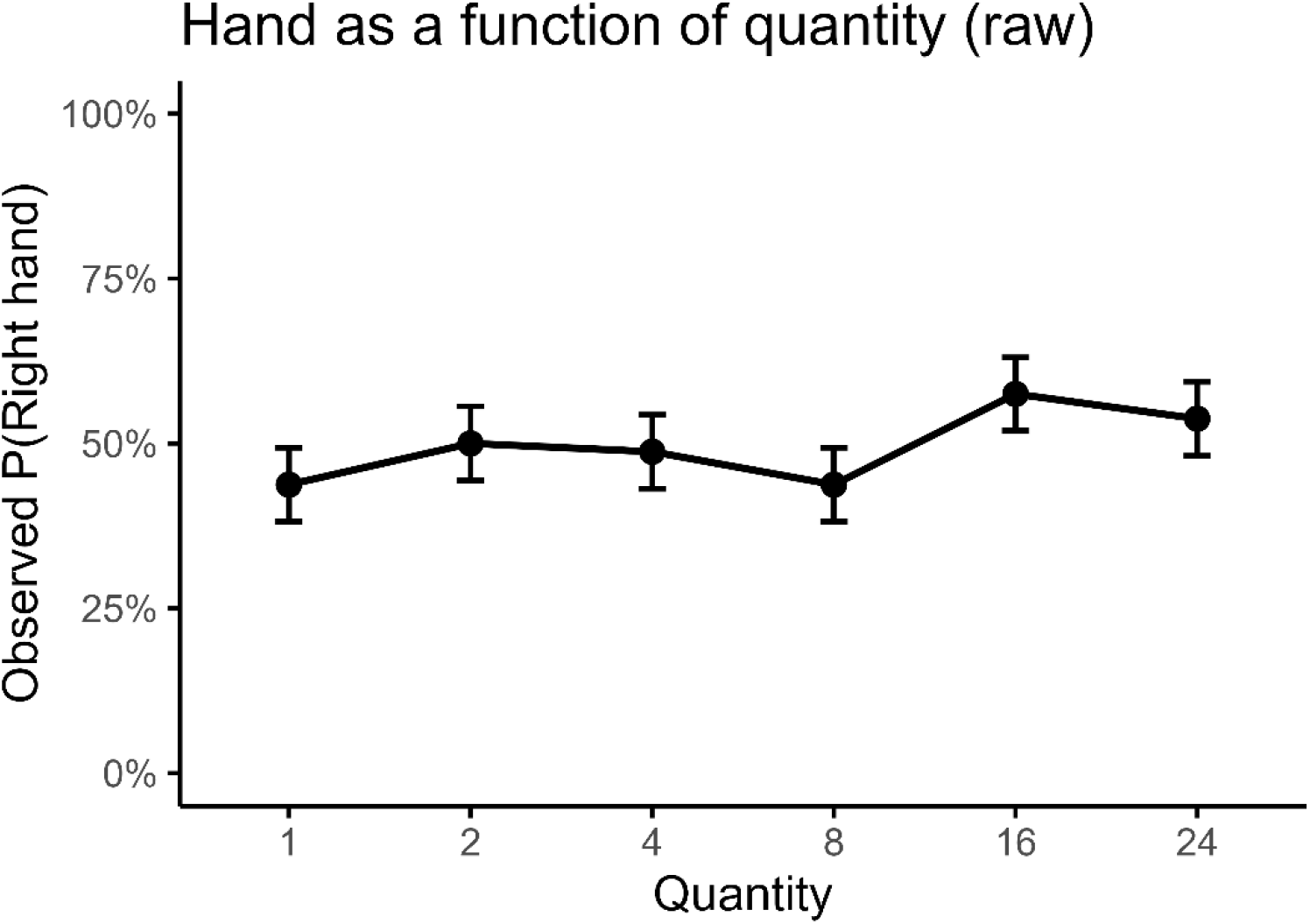
Hand preference across numerical amounts (raw proportions). Mean proportion of right-side hand use across the six numerical amounts (1, 2, 4, 8, 16, 24 items). Dots represent group means and vertical bars indicate ±1 standard error (SE). No systematic lateral bias as a function of quantity was observed.

### Statistical Analysis of Hand as a Function of Numerical Magnitude (Logarithmic Linear Trend Model)

To assess whether numerical magnitude influenced hand preference, we fitted a generalized linear mixed-effects model including a continuous numerical predictor reflecting the logarithmic structure of the quantities (log2-transformed and centered). The model included a random intercept for subject identity to account for repeated measures.

#### Results

A likelihood ratio test comparing the full model with a reduced model excluding numerical magnitude revealed a significant improvement in model fit (Δχ^2^(1) = 3.16, p* = .038; Table S8b), indicating that numerical magnitude contributed to explaining variation in hand use. Consistently, the numerical predictor was significant in the full model, showing a positive linear relationship between log-transformed quantity and right-hand use (χ^2^(1) = 4.45, p* = .034; Table S8a). This result indicates that the probability of using the right hand increased monotonically as numerical magnitude increased along a logarithmic scale (Fig. S2).

Importantly, this effect emerged in the absence of a corresponding spatial choice bias, suggesting that numerical magnitude selectively modulated response-related motor processes rather than allocentric spatial choice.

#### Statistical Analysis of Hand Preference Based on Quantity Grouping

To further examine whether hand preference differed as a function of overall numerical magnitude, we conducted an additional GLMM in which quantities were grouped into two categories: small amounts (1–8 items) and large amounts (16–24 items). The dependent variable was hand use (0 = left, 1 = right), and Amount_group (Small/Large) was entered as a fixed effect, with a random intercept for Subject ID.

#### Results

The model revealed a significant effect of Amount_group (χ^2^(1) = 7.338, p** = .007; Table S9), indicating that monkeys were more likely to use the right hand when evaluating larger numerical quantities. Model-based estimates confirmed this pattern, with the probability of right-hand use increasing from 0.413 (SE = 0.174) for small amounts to 0.599 (SE = 0.177) for large amounts. Pairwise contrasts showed that the odds of using the right hand were approximately twice as high for large quantities compared to small ones (OR = 2.12, z = 2.709, p** = .0068; Fig. 5 and Fig. S10a–b).

**Figure 5.**
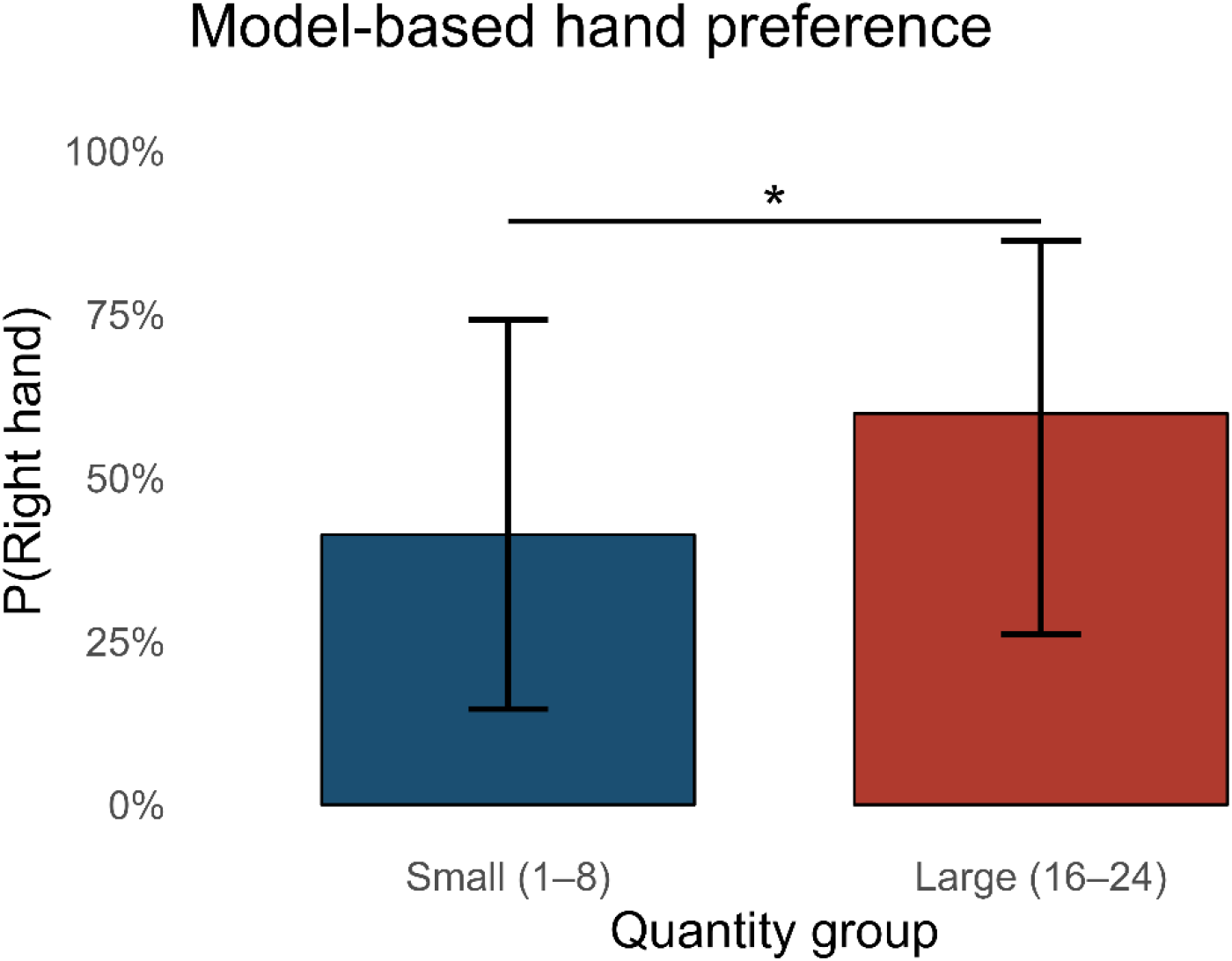
Model-based comparison of hand preference for grouped amounts. Predicted probability of right-hand use derived from the GLMM for low (1–8) vs. high (16–24) numerical amounts. Bars represent model-estimated probabilities and vertical bars indicate 95% confidence intervals. Larger quantities elicited higher right-hand use, confirming a categorical magnitude-dependent effect.

Together with the logarithmic trend analysis, these results indicate a robust magnitude-dependent modulation of hand use, supporting the view that numerical magnitude engages lateralized motor response systems even when no spatial choice bias is observed.

## Conclusions

The results indicate that absolute numerical magnitude did not generate a left–right spatial bias in external choice space. No systematic association was observed between numerical quantity and spatial choice when quantities were considered individually, along a logarithmic continuum, or when grouped into small and large values. In contrast, numerical effects emerged selectively in the motor domain: hand-use asymmetries increased for larger numerical magnitudes, particularly when quantities were treated as coarse categories or along a logarithmic scale. This dissociation suggests that numerical magnitude influenced response-related motor processes rather than allocentric spatial choice per se. This pattern is consistent with proposals that number–space interactions are partly grounded in action and manual response dynamics (Beran et al., 2019), as observed in human SNARC-like effects, and with evidence that macaques may express numerical–spatial associations preferentially in motor or relative numerical contexts rather than through direct mappings onto external space. To directly test this possibility, in Experiment 2 we made use of a habituation–dishabituation paradigm based on relative numerical changes.

### Experiment 2

#### Subjects and apparatus

In the second experiment we tested 12 individuals: 2 rhesus macaques (*Macaca mulatta*) and 10 crab-eating macaque (*Macaca fascicularis*) who participated to the Experiment 1. The age of the monkeys ranged from 3 to 14 years (mean age = 5.33, sd = 3.938). The subjects were tested in the same testing cages as experiment I and we employed the same food boards. All housing and procedures used in this research complied with the current guidelines in the matter of care and use of laboratory animals (European Community Council Directive No. 86–609), and were authorized by our local ethics board (03.10.18) and the French Ministry of Research (10.10.18; authorization no. 15091_2018071014483295_v2).

#### Experimental Procedure

In Experiment 2, we employed a structured habituation–dishabituation paradigm to examine whether spatial biases are sensitive to variations in numerical quantity. During the habituation phase, macaques were repeatedly exposed to a fixed reference numerosity, establishing a baseline representation of numerical magnitude. In the subsequent dishabituation phase, they were presented with a numerosity either larger or smaller than this reference. This design allowed us to assess whether monkeys exhibit a relative spatial bias that shifts according to numerical variation, rather than simply responding to absolute quantities. To implement this, we used two pair-wise combinations of food quantities: 4×4 (small quantity) and 16×16 (large quantity). For each subject, the experimental session consisted of two sequences (“Sequence A - Habituation large” and “Sequence B - Habituation small”) of thirty-three trials each. Both sequences were characterized by three blocks, each one consisting of a habituation phase of 10 trials with the same pairwise combination (4×4 in “habituation small” and 16×16 in “habituation large” respectively), followed by one novel trial with the other pairwise of food (novel phase: 16×16 for the “habituation small” and 4×4 for the “habituation large” respectively). Therefore, the sequence “Habituation large” consisted of thirty habituation trials with the pairwise combination 16×16 and three novel trials with 4×4. In contrast, the Sequence “Habituation small” involved thirty habituation trials with 4×4 pairwise combination and three novel trial with 16×16.

### Data Elaboration

For each animal, the experimental session was coded as previously explained for experiment

1. A second researcher coded a random 20% of videos to assess reliability. The reliability scores between coders were very high (all k > 0.8). Two individuals (*Macaca fascicularis*) did not complete the task due to technical issues during its execution. The two pairwise combinations of identical food quantities (4×4 and 16×16) were classified as follows: the 4×4 combination was considered a “small” quantity, while the 16×16 combination was considered a “large” quantity. All analyses were conducted on the last habituation trials of each block and on the dishabituation trials.

## Choice Preference Analysis

### Statistical Analysis of Lateral Choice in the Habituation/Dishabituation Paradigm

We investigated whether monkeys’ spatial choices (left vs. right) were influenced by numerical magnitude during habituation and dishabituation sequences using a generalized linear mixed model (GLMM) with a binomial distribution and a logit link function. The dependent variable was Choice (coded as 0 = left, 1 = right). Prior to fitting the final model, we tested the potential effects of Trial Quality and Subject Position in separate GLMMs. Trial Quality did not significantly influence lateral choices (χ^2^(2) = 0.805, *p* = .669), while Position (χ^2^(2) = 12.11, *p*\*\* = .002) showed significant effects (Table S11). Therefore Position was included as covariate in the final model.

The primary fixed effects of interest in the final model were Pair (numerical identity of the pair: 4 vs. 16), TrialType (Habituation or Dishabituation), and their interaction. A random intercept for Subject ID was included to account for repeated measures within individuals. Model fitting was conducted in R version 4.3.1 using RStudio 2025.05.0 Build 496. The model was estimated with the lme4 package (Bates et al., 2015), and Type III Wald chi-square tests were used to assess the significance of fixed effects via the *car* package (Fox & Weisberg, 2018). Estimated marginal means and pairwise comparisons were calculated using the *emmeans* package (Lenth, 2023).

The final model revealed a significant main effect of Pair × TrialType interaction (χ^2^(1) = 4.63, *p\** = .031), suggesting that the effect of numerical magnitude on spatial choice was modulated by trial type (habituation vs. dishabituation). Position (χ^2^(2) = 10.64, *p*\*\* = .004) also significantly predicted lateral choice. No significant main effect of Pair alone (6 or 14) (χ^2^(1) = 0.06, *p* = .809) or TrialType was found (χ^2^(1) = 2.741, *p* = .097 ; Table S12).

Model-based predictions (EMMEANS) revealed a clear TrialType × Pair interaction effect in choice behaviour. During habituation, monkeys showed no difference between the two numerical quantities (Pair 4 vs 16: OR = 1.15, p = 0.81 ;Table S14a). However, during dishabituation, their responses diverged markedly: after a numerical increase (4 → 16), monkeys were more likely to choose the right side (P = 0.676), whereas after a numerical decrease (16 → 4), they showed a lower probability of choosing right (P = 0.258) (TableS13).

This difference during dishabituation was statistically significant (Pair 4 vs 16: OR = 0.17, SE = 0.11, z = −2.67, p** = 0.007), confirming that monkeys responded asymmetrically depending on whether the number increased or decreased (Table S14).

Taken together, these results indicate that a change in numerical magnitude induced a directional spatial bias, consistent with a number–space mapping effect: decreases in quantity biased choices toward the left side of space, whereas increases biased choices toward the right (Fig. 6)

**Figure 6.**
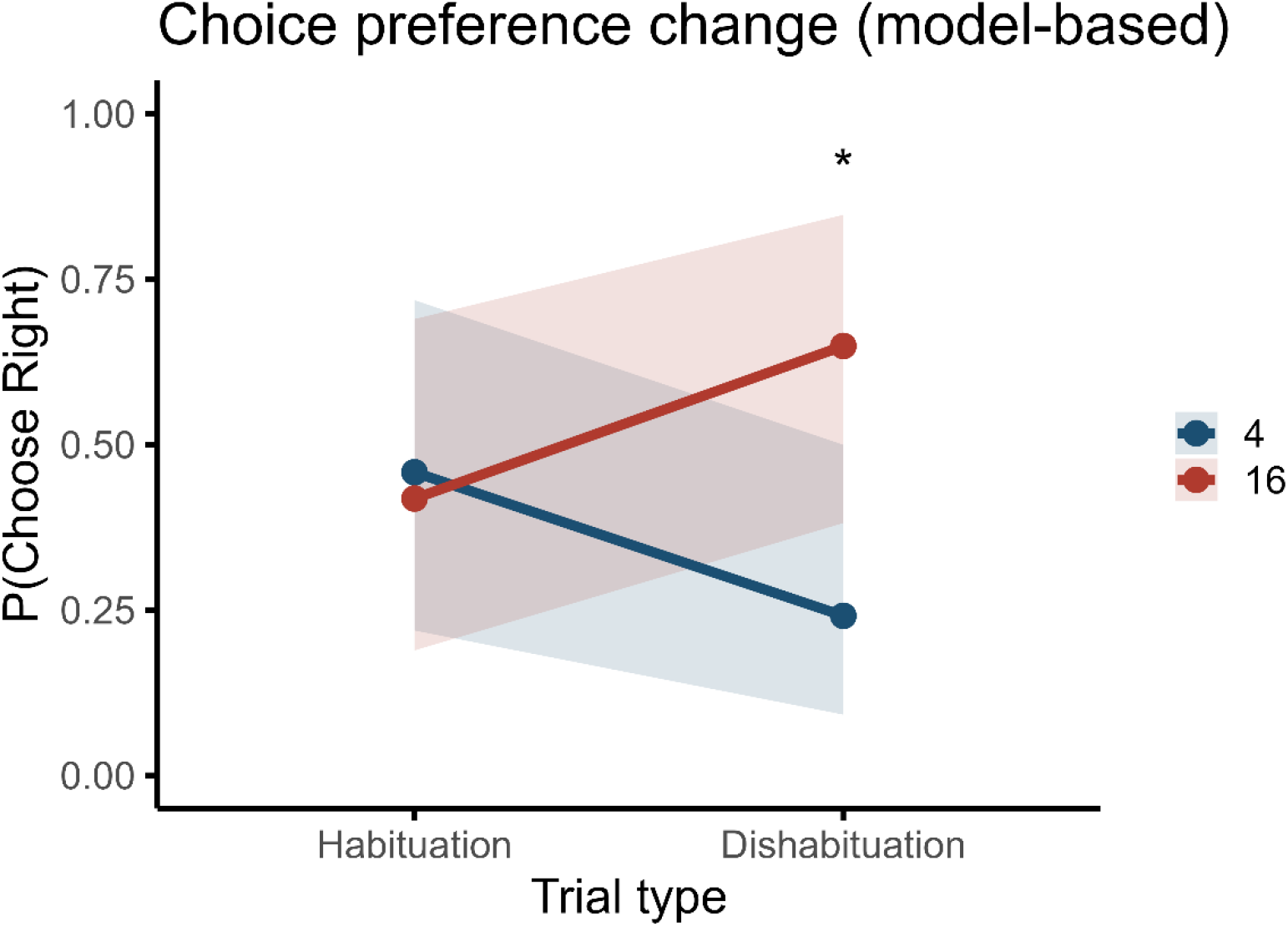
Model-based choice preference across trial types Predicted probability of choosing the right side during habituation (0) and dishabituation (1), plotted separately for Pair 4 and Pair 16. Shaded bands show 95% confidence intervals. Choice behaviour diverged specifically during dishabituation: following a numerical decrease (16→4; dishabituation Pair 4) monkeys showed fewer rightward choices, whereas following a numerical increase (4→16; dishabituation Pair 16) they showed more rightward choices, consistent with a direction-dependent number–space mapping.

### Statistical Analysis – Hand Use in Habituation/Dishabituation Paradigm

#### Statistical Analysis of Hand Use in the Habituation–Dishabituation Task

We investigated whether hand choice (Hand, coded as 0 = left, 1 = right) was influenced by numerical context during a habituation–dishabituation paradigm composed of 33 trials per block (10 habituation trials followed by 1 surprise trial, repeated three times). The main predictors were Pair (4 vs. 16), TrialType (Habituation/Dishabituation) and their interaction.

Before fitting the final model, we tested whether Trial Quality and Subject Position significantly affected hand choice using separate generalized linear mixed models (GLMMs) with a random intercept for each subject. None of these variables reached statistical significance and were therefore not included in the final model (Table S15).

The final GLMM included the interaction between Pair and TrialType as fixed effect, with a random intercept for Subject to account for repeated measures. Model fitting was performed in R version 4.3.1 using RStudio 2025.05.0 Build 496. The model was estimated using the `lme4` package (Bates et al., 2015), and fixed effects were evaluated using Type III Wald chi-square tests via the `car` package (Fox & Weisberg, 2018). Post-hoc comparisons were conducted using estimated marginal means with the `emmeans` package.

Hand preference was significantly influenced by the interaction between numerical magnitude (Pair) and Trial Type (Habituation/Dishabituation). Type III Wald χ^2^ tests revealed a significant main effect of TrialType (χ^2^(1) = 5.62, p* = .018), indicating that hand use varied between habituation and dishabituation trials. Furthermore, a significant interaction between Pair and TrialType was observed (χ^2^(1) = 10.42, p** = .0012), suggesting that the effect of numerical magnitude on hand preference was modulated by trial type. No significant main effect of Pair was found on its own (χ^2^(1) = 1.21, p = 0.27), nor was the intercept significant (χ^2^(1) = 0.688, p = 0.407; Table S16).

Estimated marginal means indicated that hand use differed as a function of numerical change during dishabituation. Specifically, when a smaller quantity followed habituation to a larger one (16 → 4), monkeys showed a marked reduction in right-hand use, whereas the opposite pattern was observed when a larger quantity followed a smaller one (4 → 16) (Table S17).

Post-hoc contrasts confirmed that this difference during dishabituation was statistically significant (Pair 4 vs Pair 16: OR = 0.078, z = −3.43, p*** = .0006; Table S18). These results indicate that hand preference differed between numerical pairs specifically during dishabituation trials.

Overall, these findings indicate that numerical magnitude modulates hand use in a direction-specific manner, revealing a spatial–numerical association expressed at the level of motor responses during numerical expectancy violations (Fig.7).

**Figure 7.**
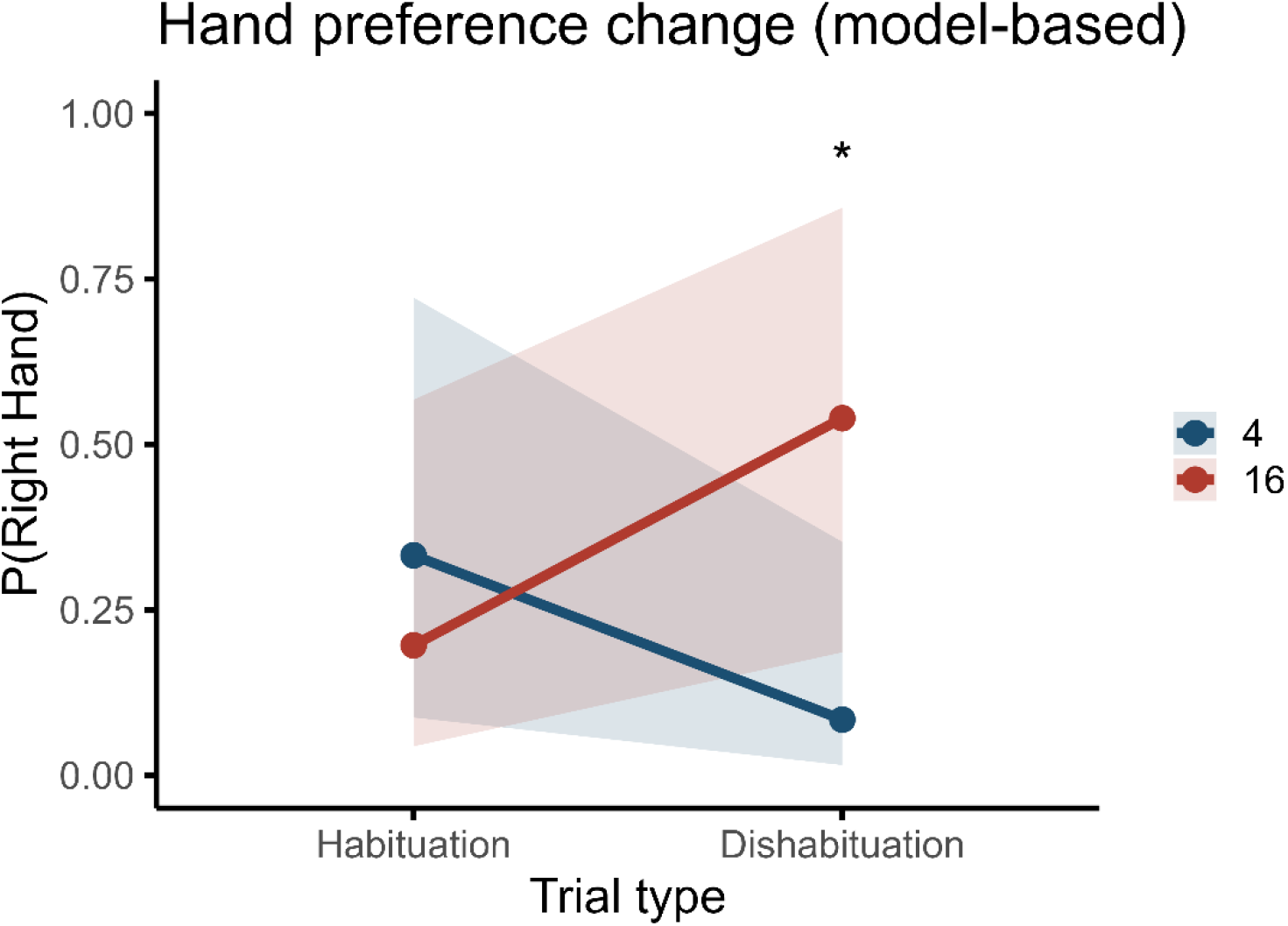
Model-based hand preference across trial types Predicted probability of right-hand use during habituation and dishabituation, plotted separately for Pair 4 and Pair 16. Shaded bands show 95% confidence intervals (back-transformed from the logit scale). Hand use diverged during dishabituation: right-hand use was markedly reduced when the dishabituation numerosity was small (16→4), whereas it was higher when the dishabituation numerosity was large (4→16), indicating a direction-dependent modulation of manual responses.

## Conclusions

Experiment 2 shows that macaques’ lateralized responses are sensitive to relative numerical change when magnitude is evaluated against a habituated reference. Rather than exhibiting a fixed left–right mapping for absolute quantities, the animals’ spatial choices varied systematically depending on whether the numerical value increased or decreased relative to the habituation baseline. Specifically, numerical increases were associated with a shift toward rightward responses, whereas numerical decreases were associated with a relative reduction in rightward choices. This direction-dependent modulation emerged primarily during dishabituation trials, suggesting that lateralized spatial responses are particularly engaged when an established expectation is violated.

A similar, and even clearer, pattern was observed in hand use. Manual responses diverged sharply depending on whether the numerical change involved an increase or a decrease, with right-hand use being reduced following decreases and enhanced following increases. This indicates that number-related lateralization may be expressed most strongly at the level of motor output, especially in situations that require active comparison with a previously learned reference.

Taken together, these findings suggest that numerical lateralization in macaques does not reflect a rigid spatial mapping of quantity. Instead, it appears to arise dynamically in contexts that involve expectancy formation and comparison processes. Such effects may be mediated by attentional engagement, motivational salience, or motor planning mechanisms that become particularly relevant when numerical changes violate established predictions. Further work will be necessary to disentangle these potential mechanisms and clarify how reference-based magnitude processing interacts with hemispheric asymmetries in non-human primates.

## Discussion

The present study aimed at assessing the occurrence of a left-to-right Mental Number Line (MNL) in two species of macaques using spontaneous food-related tasks. In Experiment 1, we investigated whether macaques exhibit an oriented mental number line in a spontaneous equal–equal comparison task, in which two identical numerical quantities were presented laterally. In contrast, Experiment 2 explored the emergence of spatial–numerical associations when a change in numerosity occurred relative to a habituated reference, requiring monkeys to rely on comparative rather than absolute numerical evaluation.

Across experiments, our results indicate that a directional number–space mapping does not reliably emerge when numerical quantities are evaluated in isolation but becomes evident when magnitude is processed relative to an established reference. In Experiment 2, we observed a clear direction-dependent modulation of spatial choice during dishabituation trials. Specifically, following a numerical increase (4 → 16), monkeys showed a higher probability of rightward choices, whereas following a numerical decrease (16 → 4), they showed a reduced probability of choosing the right side. This pattern indicates that the direction of numerical change, rather than absolute magnitude per se, is critical for eliciting spatial biases.

This finding is consistent with the idea that repeated exposure to a reference numerosity shifts the internal baseline used for numerical evaluation (Vallortigara, 2018). After habituation to a large quantity, smaller numerosities may be perceived as relatively more “leftward,” whereas the opposite shift occurs following habituation to smaller values.

In contrast, Experiment 1 did not provide evidence for a stable left–right mapping based on absolute numerical magnitude. Spatial choice was robustly influenced by non-numerical factors such as subject position, but neither numerical identity, numerical magnitude treated as a continuum, nor coarse magnitude grouping reliably modulated lateral choice. Together, these findings suggest that relative numerical evaluation, rather than absolute magnitude, is a key condition for the emergence of spatial–numerical associations in macaques, consistent with proposals emphasizing the importance of comparison processes in SNARC-like effects (Vallortigara, 2018 ; Vallortigara, 2021).

The design of Experiment 2 closely parallels paradigms previously used in other species, including chicks (Rugani et al., 2015), honeybees (Giurfa et al., 2002), and human newborns (Di Giorgio et al., 2019), all of which reported directional number–space mappings following habituation. Our findings align with this body of work but differ from a more recent study in rhesus macaques (Beran et al., 2019), which did not observe comparable spatial–numerical effects in an equal-quantity touchscreen task. This discrepancy may reflect differences in task structure, sample size, and prior experience with numerical tasks. Unlike the trained monkeys tested by Beran and collegues, our subjects were numerically naïve, and their responses may therefore reflect more spontaneous numerical processing, less influenced by top-down cognitive control mechanisms.

Another critical difference concerns stimulus ecology. Whereas Beran and colleugues employed digital stimuli with constant reward independent of numerical evaluation, our experiments relied on biologically relevant food stimuli and direct monitoring of visual attention. Previous work has shown that ecological relevance can enhance motivation and engagement in numerical tasks (Addessi et al., 2008), and that contextual factors such as stimulus format and trial structure can substantially modulate the expression of SNARC-like effects (Basso Moro et al., 2018).

Interestingly, a partially dissociable pattern emerged for hand use compared to spatial choice. In Experiment 1, hand preference did not differ across the six exact Pair levels in the categorical model, but exploratory analyses converged in indicating a magnitude-related motor bias: both the log-scaled predictor and the small/large grouping showed increased right-hand use for larger quantities. This pattern suggests that numerical magnitude can modulate response selection at the motor level even when it does not reliably bias allocentric left–right choice.

In Experiment 2, manual responses were even more clearly shaped by relative numerical change during dishabituation, consistent with a direction-dependent modulation of action (decreases associated with reduced right-hand use; increases associated with increased right-hand use). Together, these findings suggest that number–space interactions in macaques are most robustly expressed through lateralized motor outputs, rather than as fully abstract spatial preferences, and that only under specific comparative contexts do these interactions extend to overt spatial choice.

Taken together, the results of Experiment 2, in combination with previous findings in human infants and nonhuman animals, lend support to the idea that number–space associations may rely on an early-developing, potentially innate component of numerical representation. In particular, the observation that numerical magnitude modulated action selection and lateralized motor responses under comparative conditions suggests that such associations can emerge in the absence of explicit instructions or culturally acquired spatial conventions.

Importantly, the present findings leave open the possibility that number–space associations, while grounded in early-emerging constraints, may subsequently be refined or reorganized through experience (see Introduction). In nonhuman primates, experiences may arise from sensorimotor contingencies and task structure, whereas in humans they may be further canalized by culturally specific practices such as reading and writing. In this sense, early-emerging numerical biases provide a substrate upon which different species-specific patterns of number–space mapping can be constructed.

Crucially, evidence for number–space associations in infants and nonhuman animals relies on methodologies that differ substantially from those typically employed in adult human studies. Consistent with this view, recent work has suggested that SNARC-like effects are reliably observed in implicit tasks even in traditional societies without written culture (Eccher et al., 2023).

Overall, the present findings support the existence of a left-to-right numerical bias in macaques that emerges most reliably in contexts involving expectancy violations in the estimation of magnititudes.

