## Supplementary material for "Number-Space Association in Macaques": Supplemetary Materials

Supplementary Tables report the full statistical output of the generalized linear mixed models (GLMMs) used in Experiments 1 and 2. For clarity and consistency across analyses, all tables report predictors, Wald $\chi^2$ statistics, degrees of freedom (df), and associated p-values.

### *Experiment 1 – Choice Preference Analysis*

Table S1. Type III Wald Chi-square Tests – Control Models (Response: Choice)

| **Control Variable** | **Predictor** | **$\chi^2$** | **df** | **p-value** | **Significance** |
|---|---|---|---|---|---|
| Quality | (Intercept) | 0.600 | 1 | 0.438 | |
| | Quality | 4.35 | 2 | 0.114 | |
| Position | (Intercept) | 7.560 | 1 | 0.005 | ** |
| | Position | 25.29 | 2 | < 0.001 | *** |

Table S2. Type III Wald Chi-square Tests for Final GLMM (Response: Choice)

| **Predictor** | **$\chi^2$** | **Df** | **p-value** | **Significance** |
|---|---|---|---|---|
| (Intercept) | 9.439 | 1 | 0.002 | ** |
| Pair | 6.44 | 5 | 0.266 | |
| Position | 22.72 | 2 | < 0.001 | *** |

Table S3. Type III Wald Chi-square Tests – Logarithmic Linear Trend Model (Response: Choice)

| **Predictor** | **Effect** | **$\chi^2$** | **df** | **p-value** | **Significance** |
|---|---|---|---|---|---|
| Intercept | (Intercept) | 7.719 | 1 | 0.005 | ** |
| Pair_log2_c | Logarithmic linear trend (centered) | 2.085 | 1 | 0.149 | |
| Position | Position | 25.35 | 2 | < 0.001 | *** |

Table S4. Likelihood-Ratio Test for Logarithmic Linear Trend Model

| **Model comparison** | **$\Delta\chi^2$** | **df** | **p-value** | **Significance** |
|---|---|---|---|---|
| Full model vs. Null model | 2.07 | 1 | 0.151 | — |

Table S5. Type III Wald Chi-square Tests – Grouped Quantity Model (Response: Choice)

| **Predictor** | **$\chi^2$** | **df** | **p-value** | **Significance** |
|---|---|---|---|---|
| (Intercept) | 8.446 | 1 | 0.003 | ** |
| Amount_group | 2.263 | 1 | 0.132 | |

| Predictor | $\chi^2$ | df | p-value | Significance |
|---|---|---|---|---|
| Position | 24.055 | 2 | < 0.001 | *** |

### *Experiment 1 – Hand Preference Analysis*

Table S6. Type III Wald Chi-square Tests – Control Models (Response: Hand)

| Control Variable | Predictor | $\chi^2$ | Df | p-value | Significance |
|---|---|---|---|---|---|
| Quality | (Intercept) | 0.429 | 1 | 0.512 | |
| | Quality | 1.610 | 2 | 0.447 | |
| Position | (Intercept) | 0.991 | 1 | 0.319 | |
| | Position | 5.090 | 2 | 0.078 | |

Table S7. Type III Wald Chi-square Tests for Final GLMM (Response: Hand)

| Predictor | $\chi^2$ | df | p-value | Significance |
|---|---|---|---|---|
| (Intercept) | 0.579 | 1 | 0.447 | |
| Pair | 9.903 | 5 | 0.078 | |

Table S8a. Type III Wald Chi-square Tests – Logarithmic Linear Trend Model (Response: Hand)

| Model Term | Predictor | $\chi^2$ | df | p-value | Significance |
|---|---|---|---|---|---|
| Logarithmic Linear Trend | (Intercept) | 0.019 | 1 | 0.889 | |
| | Pair_log2_c | 4.452 | 1 | 0.034 | * |

Table S8b. Likelihood-Ratio Test for Logarithmic Linear Trend Model

| Model | npar | AIC | BIC | logLik | $\chi^2$ | df | p-value | Significance |
|---|---|---|---|---|---|---|---|---|
| Trend null model | 2 | 420.85 | 429.20 | -208.43 | — | — | — | — |
| Full trend model | 3 | 418.54 | 431.06 | -206.27 | 4.32 | 1 | 0.038 | * |

Table S9. Type III Wald Chi-square Tests – – Grouped Quantity Hand Model

| Predictor | $\chi^2$ | Df | p-value | Significance |
|---|---|---|---|---|
| (Intercept) | 0.239 | 1 | 0.625 | |
| Amount_group | 7.338 | 1 | 0.007 | ** |

Table S10a. Estimated Marginal Means and Contrast – Grouped Quantity Hand Model

| Amount Group | Prob(Right Hand) | SE | 95% CI |
|---|---|---|---|
| Small (1–8) | 0.413 | 0.174 | [0.147, 0.742] |
| Large (16–24) | 0.599 | 0.177 | [0.261, 0.863] |

Table S10b. Pairwise Contrast

| Contrast (Reference) | Odds Ratio | SE | z | p-value |
|---|---|---|---|---|
| Large (16–24) vs Small (1–8) | **2.12** | 0.277 | 2.709 | **0.0068** |

### *Experiment 2 – Choice Preference Analysis*

Table S11. Control Models – Preliminary GLMMs for Choice (Habituation/Dishabituation Paradigm)

| Control variable | Predictor | $\chi^2$ | Df | p-value |
|---|---|---|---|---|
| Quality | Intercept | 0.699 | 1 | 0.403 |
| | Quality | 0.805 | 2 | 0.669 |
| Position | Intercept | 4.495 | 1 | 0.034* |
| | Position | 12.115 | 2 | 0.002 ** |

Table S12. Type III Wald chi-square tests for the final GLMM (Choice as response)

| Predictor | $\chi^2$ | df | p-value |
|---|---|---|---|
| (Intercept) | 2.432 | 1 | 0.119 |
| Pair | 0.059 | 1 | 0.809 |
| TrialType | 2.741 | 1 | 0.097 |
| Position | 10.645 | 2 | 0.0048 ** |
| Pair × TrialType | 4.636 | 1 | 0.0313 * |

Table S13. Estimated marginal means for choice responses as a function of TrialType (Habituation/Dishabituation) and numerical Pair (4 vs 16).

| TrialType | Pair | prob | SE | 95% CI lower | 95% CI upper |
|---|---|---|---|---|---|
| Habituation | 4 | 0.504 | 0.158 | 0.228 | 0.778 |
| Dishabituation | 4 | 0.258 | 0.126 | 0.087 | 0.559 |
| Habituation | 16 | 0.468 | 0.160 | 0.199 | 0.757 |

| | | | | | |
|---|---|---|---|---|---|
| Dishabituation | 16 | 0.676 | 0.138 | 0.371 | 0.878 |

Table S14. Pairwise contrasts (within each TrialType: Habituation/Dishabituation condition)

| TrialType | Contrast | Odds ratio | SE | z | p |
|---|---|---|---|---|---|
| Habituation | Pair4 / Pair16 | 1.158 | 0.705 | 0.241 | 0.809 |
| Dishabituation | Pair4 / Pair16 | 0.166 | 0.111 | −2.673 | **0.007** |

### *Experiment 2 – Hand Preference Analysis*

Table S15. Control Models – Preliminary GLMMs for Hand

| Control Variable | Predictor | $\chi^2$ | df | p-value |
|---|---|---|---|---|
| Quality | (Intercept) | 0.019 | 1 | 0.888 |
| | Quality | 0.369 | 2 | 0.831 |
| Position | (Intercept) | 2.874 | 1 | 0.090 |
| | Position | 0.956 | 2 | 0.620 |

Table S16. Type III Wald $\chi^2$ tests for final model predicting hand use

| Predictor | $\chi^2$ | df | p-value |
|---|---|---|---|
| (Intercept) | 0.688 | 1 | 0.407 |
| Pair | 1.208 | 1 | 0.272 |
| TrialType | 5.620 | 1 | 0.0178 * |
| Pair × TrialType | 10.417 | 1 | 0.0012 ** |

Table S17. Estimated marginal means for hand responses as a function of TrialType (Habituation/Dishabituation) and numerical Pair (4 vs 16).

| TrialType | Pair | prob | SE | 95% CI lower | 95% CI upper |
|---|---|---|---|---|---|
| Habituation | 4 | 0.332 | 0.187 | 0.087 | 0.722 |
| Dishabituation | 4 | 0.084 | 0.069 | 0.015 | 0.353 |
| Habituation | 16 | 0.196 | 0.135 | 0.043 | 0.567 |
| Dishabituation | 16 | 0.539 | 0.208 | 0.185 | 0.858 |

Table S18. Pairwise contrasts (within each TrialType: Habituation/Dishabituation condition)

| **TrialType** | **Contrast** | **Odds ratio** | **SE** | **z** | **p** |
|---|---|---|---|---|---|
| Habituation | Pair4 / Pair16 | 2.037 | 1.319 | 1.099 | 0.2716 |
| Dishabituation | Pair4 / Pair16 | 0.078 | 0.058 | −3.432 | **0.0006** |

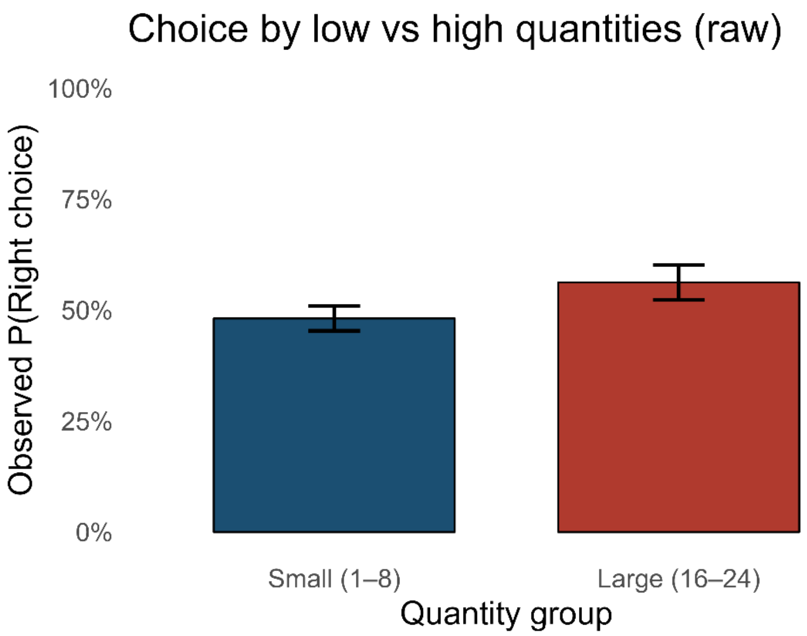


***Figure S1.*** *Choice raw grouped amounts*

*For completeness, raw proportions for grouped numerical quantities are presented, although no significant effects were detected. Bars represent observed proportions and vertical bars indicate ±1 standard error (SE).*

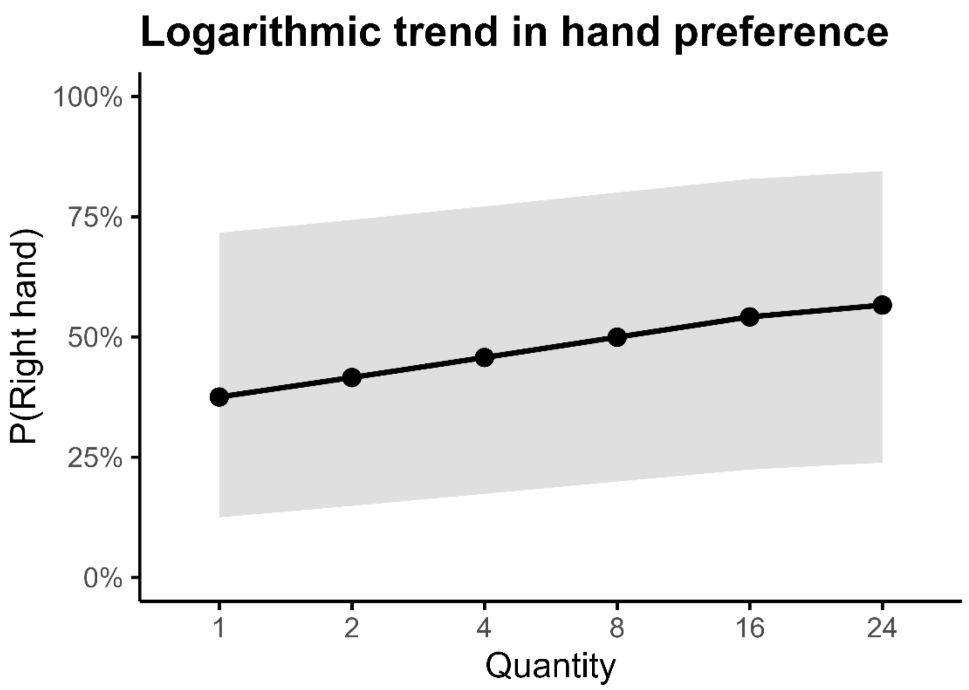


***Figure S2.*** *Model-based log2-linear trend of numerical amount on hand preference.*
*Predicted probability of right-hand use derived from the generalized linear mixed-effects model, plotted at the six experimental quantities (1, 2, 4, 8, 16, 24). Shaded areas indicate 95% confidence intervals.*
*The model revealed a significant positive logarithmic (log2-transformed) relationship between numerical magnitude and right-hand choice.*

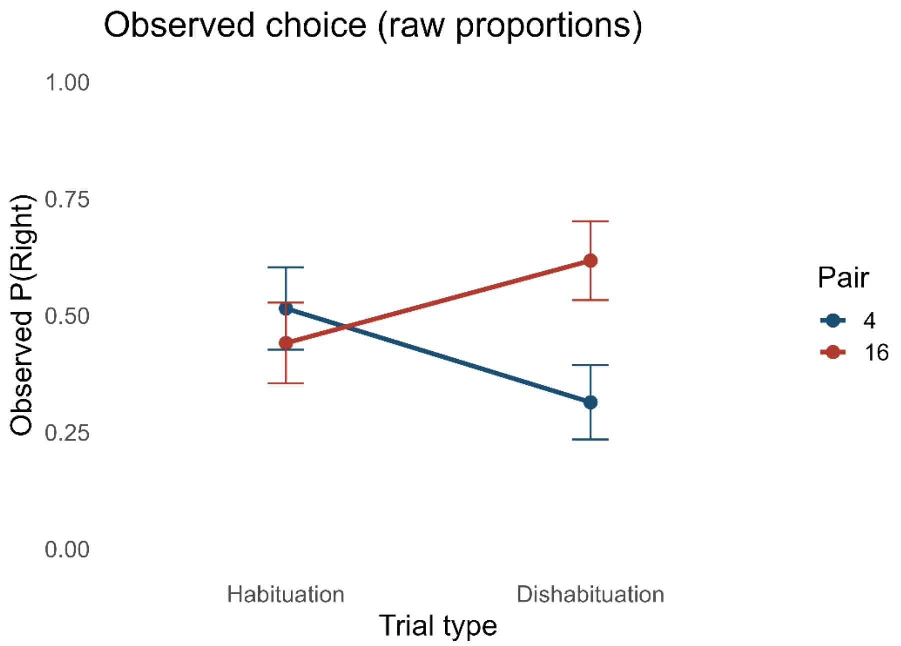

***FigureS3.*** *Observed choice proportions (raw data) across trial types.*
*Observed probability of choosing the right side (mean ±1 SE) for Pair 4 and Pair 16 during habituation and dishabituation trials (end-of-habituation trial and subsequent dishabituation trial from each block). Raw proportions mirror the model-based pattern, with divergence emerging primarily during dishabituation trials.*

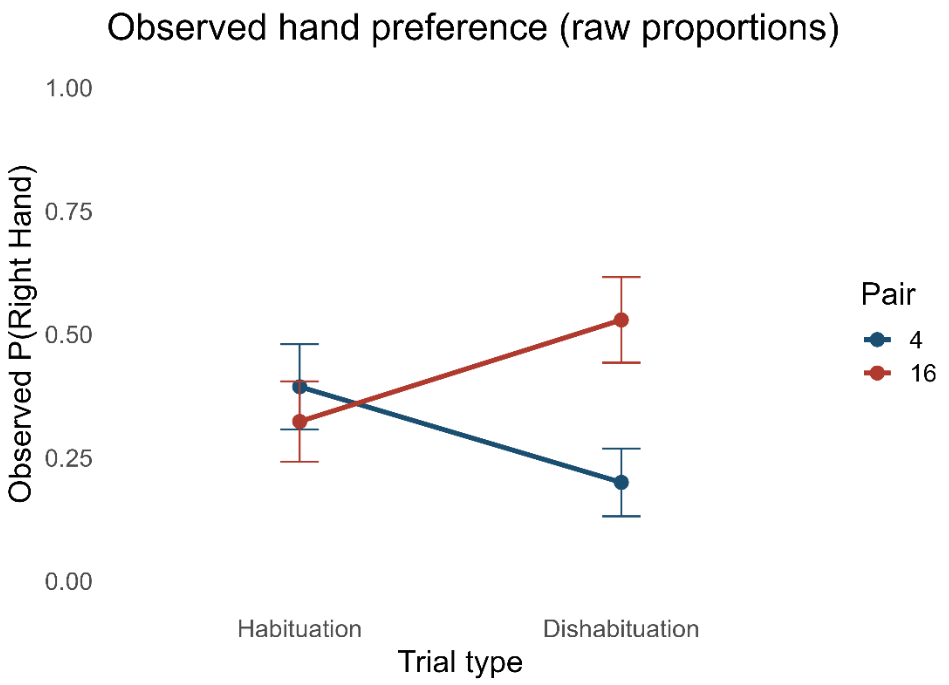


***Figure S4.*** *Observed hand-use proportions (raw data) across trial types.*
*Observed probability of using the right hand (mean ±1 SE) for Pair 4 and Pair 16 during habituation and dishabituation trials (end-of-habituation trial and subsequent dishabituation trial from each block). A separation between conditions is evident during dishabituation, consistent with the direction-dependent modulation of manual responses shown in the GLMM.*
